# Remarkably high rate of meiotic recombination in the fission yeast *Schizosaccharomyces pombe*

**DOI:** 10.1101/2022.12.12.520044

**Authors:** Qichao Lian, Laetitia Maestroni, Maxime Gaudin, Bertrand Llorente, Raphael Mercier

## Abstract

In most eukaryotes, the number of meiotic crossovers (COs) is limited to 1–3 per chromosome, which are prevented from occurring close to one another by CO interference. The fission yeast *Schizosaccharomyces pombe*, an exception to this general rule, lacks CO interference and seems to have the highest CO number per chromosome. However, global CO frequency was indirectly estimated in this species, raising doubts about this exceptional recombination level. Here, we used an innovative strategy to directly determine COs genome-wide in *S. pombe*. We confirm the absence of crossover interference and reveal the presence of co-variation in CO number across chromosomes within tetrads, suggesting that a limiting pro-CO factor varies stochastically between meiocytes. CO number per chromosome varies linearly with chromosome size, with the three chromosomes having, on average, 15.9, 12.5, and 7.0 COs, respectively. This is significantly lower than previous estimates but reinforces *S. pombe’s* exceptional status as the eukaryote with the highest CO number per chromosome described to date and among the species with the highest rate of COs per unit of DNA.

## Introduction

During meiosis, recombination is initiated by the formation of numerous programmed DNA double-strand breaks (DSBs). The repair of DSBs results in crossovers (COs), which are instances of reciprocal recombination between homologous chromosomes, and non-crossovers (NCOs), which are local tracts of non-mendelian segregation. Two main conserved pathways are responsible for CO formation. In most eukaryotes, the majority of COs, called class I COs, are specifically promoted by the group of ZMM proteins (Zip1-4, Msh4/5, and Mer3), while a minority of class II COs are dependent on other factors, including the structure-specific nuclease Mus81. Besides differences in the molecular pathways, class I COs are sensitive to CO interference, a phenomenon that prevents COs from being positioned close to each other along chromosomes, while class II COs are not subjected to interference [1–3]. The fission yeast *Schizosaccharomyces pombe* has long been used as a model to gain insights into meiotic mechanisms [4–6]. *S*. *pombe* appears to be an outlier in terms of CO control. First, CO numbers per chromosome were described to be markedly elevated, with a total genetic size of 2,250 cM for three chromosomes. This corresponds to a total of 45 COs per meiosis, and 19:11:5 COs from the largest to the smallest chromosome, respectively [6, 7]. Nineteen COs per meiosis on a single chromosome is exceptional, when considering that 80% of the analysed chromosomes in eukaryotes experience less than three [2, 8]. Second, CO interference is completely absent in *S. pombe*, with closely spaced, double COs occurring as expected if these were independent events [9, 10]. The meiotic machinery is also unusual in *S. pombe*: both synaptonemal complexes (SCs) and the class I pathway are lacking, with COs being solely dependent on the Mus81 pathway [10, 11]. However, pervasive hybrid sterility and large chromosome rearrangements [12–15] prevent the use of hybrids obtained by crossing natural isolates for accurate measurement of meiotic recombination, and a genome-wide crossover profile is still missing in *S. pombe*.

## Results and Discussion

Here, we used EMS mutagenesis to introduce a few hundred SNPs genome-wide in an isogenic strain and explored the meiotic recombination landscape in *S. pombe*. First, two independent isogenic haploid strains with the same background as the reference strain and with opposite mating types were subjected to moderate EMS treatment to introduce mutations genome-wide. Then, they were mated and sporulated to produce tetrads through meiosis (Figure 1A). To have a robust estimation of the recombination frequency, we generated and sequenced a total of 90 tetrads of three independent crosses (29, 30 and 33 tetrads, respectively, Table S1). To define the informative SNPs for each population, we deep-sequenced the parental genomes (25-91x, Table S2), and identified 161, 121 and 172 well-supported and randomly distributed mutation markers that provided good coverage of the three chromosomes (Table S3, Figure S1). The median interdistance marker is 56 kb and corresponds to the resolution at which recombination sites could be detected and assigned. Based on recombination landscapes from other species, such a resolution is suitable for crossover detection, but not for non-crossover detection that are in the kilobase range in budding yeasts [16–18].

**Figure 1.**
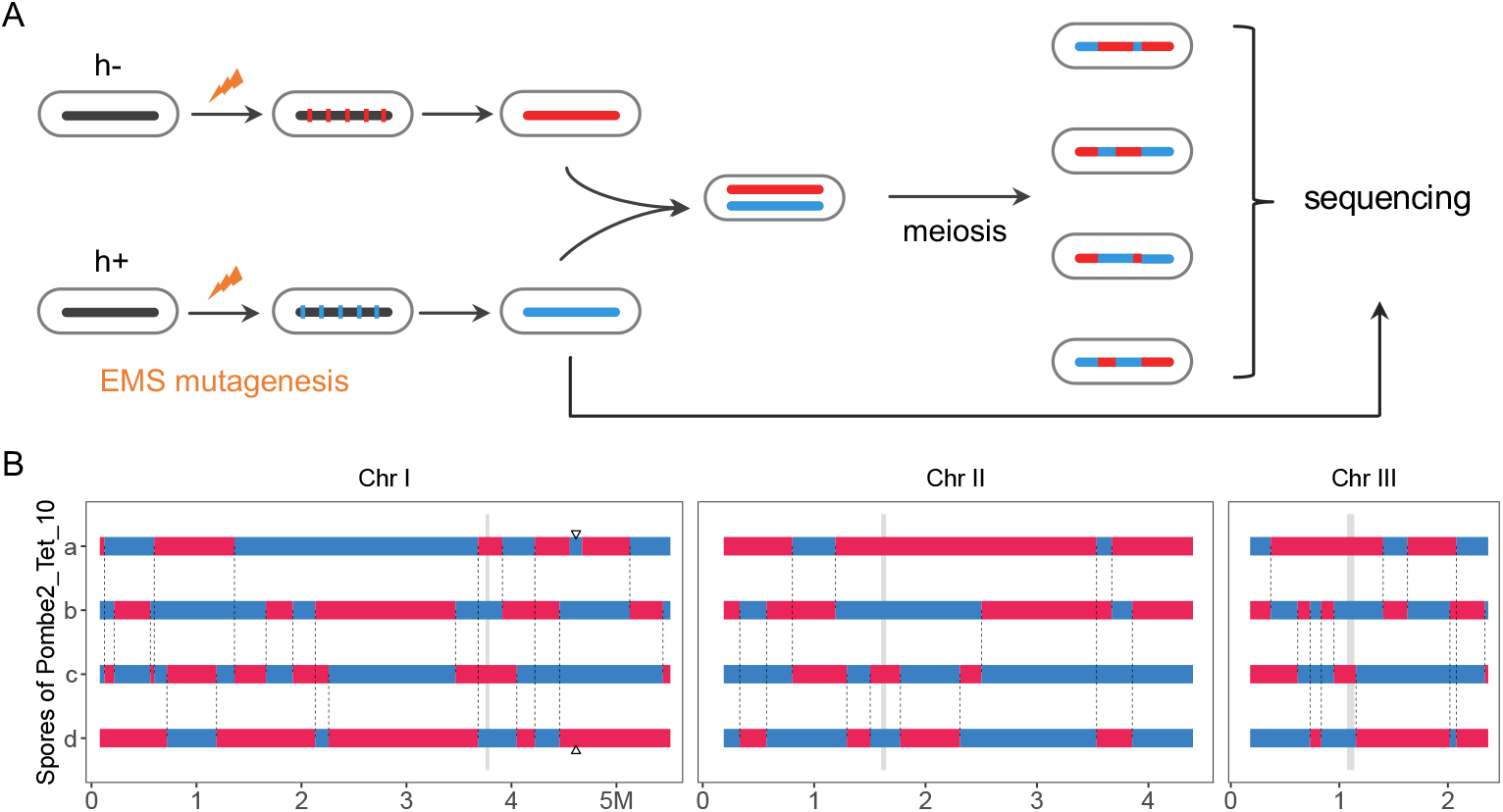
Experimental design for meiotic recombination analysis in *Schizosaccharomyces pombe* through mutagenesis of isogenic strains and tetrad sequencing. (A) SNPs (red and blue vertical bars) were introduced into the *S. pombe* genome (horizontal bar) by EMS mutagenesis in parallel in h+ and h-cells. After mutagenesis, *de novo* mutations were identified by whole genome sequencing. Cells of opposite mating type with optimal SNP distributions were mated to generate a hybrid and then tetrads after meiosis. Tetrads were dissected by micromanipulation and spores were allowed to form colonies, from which DNA was extracted and sequenced. Three independent repeats were performed. (B) CO and GC detection in a representative tetrad (Pombe2_Tet_10). The detected COs and GCs are indicated by vertical dashed lines connecting the two exchanged chromatids. A GC is indicated by a triangle. The four chromatids are color-coded according to the parental genotypes.

Using tetrad-based segregation profiles of the informative SNPs, we identified a total of 2,746 COs across the 90 meioses (median resolution 113.5 kb, Figure S2, Table S4). This corresponds to an average of 30.5 COs per tetrad (Figure 2A), and an average of 13.7, 10.8, and 6.0 COs for chromosome I, II and III, respectively (Figure 2B, 2D, Figure S3C). The distribution of the total CO number per tetrad ranged from 13 to 36 and is not significantly different from the Poisson distribution (Figure 2A). For individual chromosomes, the number of COs also follows a Poisson distribution (Figure 2D), which contrasts with many species where CO interference reduces the variance in the number of COs per chromosome [19]. Accordingly, we found that this variance of CO numbers per chromosome is larger in *S. pombe* than in *S. cerevisiae* (Figure S4). We detected at least one CO in all chromosomes in each tetrad except one where no crossover was detected on chromosome III (Figure 2D, Figure S3C). This does not imply the existence of a mechanism ensuring at least one CO per chromosome, as the elevated numbers of COs in *S. pombe* mean that the probability of an achiasmate chromosome (one without contacts with another chromosome) is ~1/1000.

**Figure 2.**
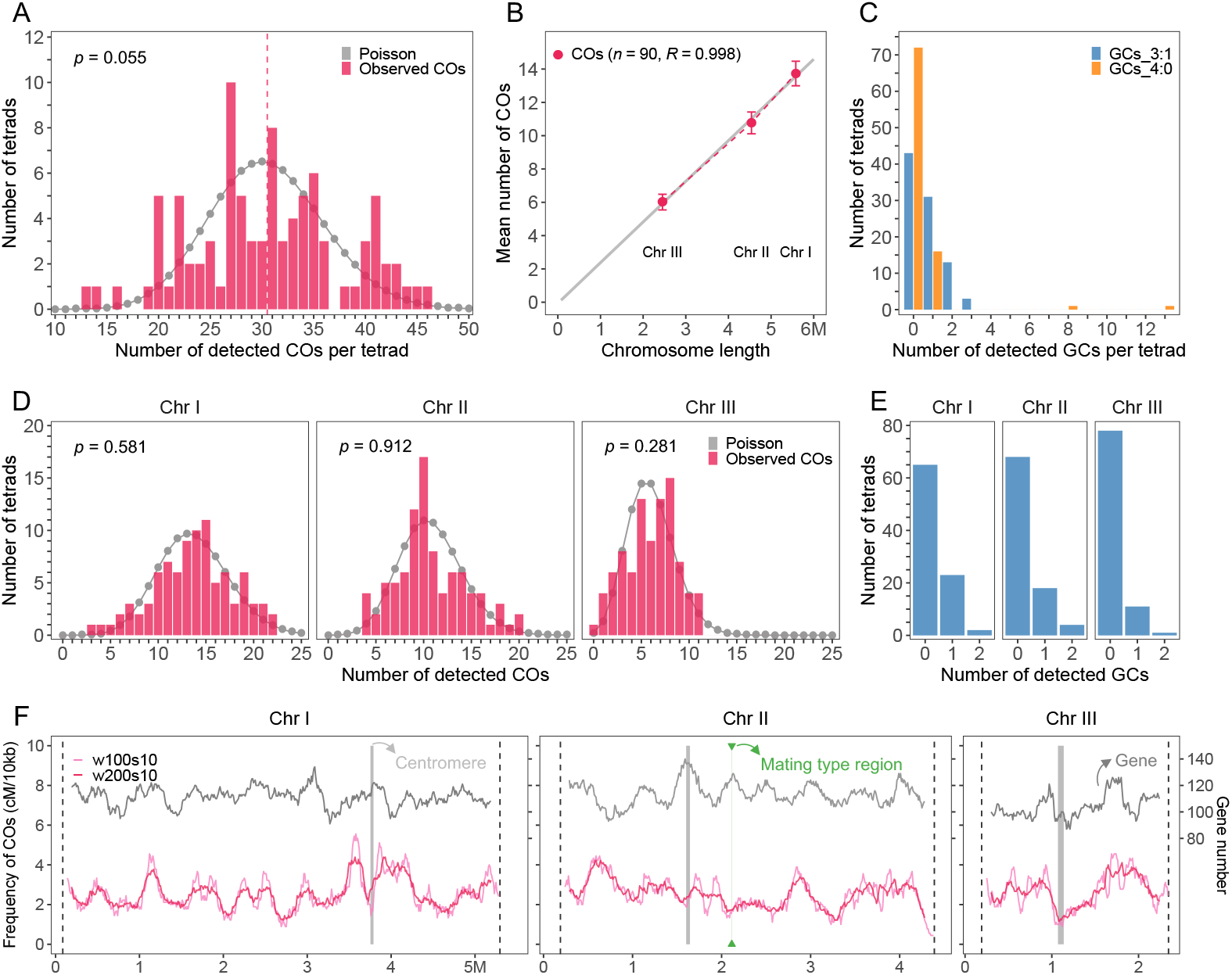
The number and distribution of COs and GCs in *S. pombe*. (A) The distribution of CO numbers per tetrad. The grey points and line represent the Poisson fitted distribution. The Mann–Whitney test was used to evaluate the differences in distributions of CO numbers between the observed and Poisson distributions. The vertical dashed line indicates the mean value. (B) Analysis of CO numbers and chromosome sizes (Mb). Crossover number and chromosome size are linearly correlated (Mb). The fitted linear regression is indicated by the grey line. The sample sizes and Pearson’s correlation coefficient are indicated in parentheses. The mean number of COs is shown with 90% confidence interval. (C) The distribution of number of GCs with 3:1 and 4:0 allelic segregation per tetrad, separately. (D) The distribution of CO numbers per chromosome and per tetrad. The grey points and line represent the Poisson fitted distribution. The Mann–Whitney test was used to evaluate the differences between observed and Poisson distributions. (E) The distribution of number of 3:1 GCs per chromosome and per tetrad. (F) The chromosomal distribution of COs with 100 kb (pink, w100s10) and 200 kb (red, w200s10) sliding window size and 10 kb step size, and genes (black) with 200 kb sliding window size and 10 kb step size, separately. The centromeric regions are indicated by grey shading. The mating type position is indicated with green shading.

Although most detected events appear as a single crossover per interval, with two chromatids showing exchange and two unmodified chromatids, we identified 108 cases of double-crossovers (DCOs) in a single interval with the four chromatids involved (Figure S5). Because of our relatively low marker density (ca. one marker per 56 kb), some DCOs may be missed in some intervals. Only DCOs involving the four chromatids of a bivalent (4C-DCO events) occurring in one interval can be detected with the flanking markers (Figure S6). However, a DCO that involves the same two chromatids (2C-DCO events) is undetectable with the flanking markers. If three chromatids are involved in a DCO event in a given interval (3C-DCO events), it is detected as a single CO (Figure S6). As chromatid interference is absent (see below), the number of two- and three-chromatid DCOs, and thus of missed DCOs, can be estimated from the number of observed four-chromatid DCOs. A total of 108 4C-DCOs were observed, leading to an estimate of 108 2C-DCOs (216 missed COs) events and 216 3C-DCOs (216 DCOs that were detected as single COs) (Table S5). We thus added 432 COs to our CO count, resulting in a total estimate of 3,178 COs in 90 meioses, corresponding to a corrected average of 35.3 COs per meiosis: 15.9, 12.5 and 7.0 COs chromosome I, II and III, separately (793 cM, 625 cM and 347 cM, respectively. Figure 3, Table S5). A previous estimate of the genetic size, based on the compilation of segregation analyses of pairs of phenotypic markers, was ~20% larger [9], likely because of the inaccuracy of this indirect approach for the total size estimate. Crossover numbers were also measured by sequencing ten spores produced by a hybrid between two closely related *S. pombe* strains, leading to an estimate of 9.6 COs per tetrad for chromosome I, 10.2 for chromosome II, and 6.8 for chromosome III [12]. This is compatible with our results, except for chromosome I where the reduced recombination frequency likely results from the presence of a large genomic inversion between these two strains that suppresses recombination events. CO numbers per chromosome are linearly correlated with the chromosome length (Pearson’s R = 0.998), with an intercept of −0.066 and an additional 2.44 COs per megabase (Figure 2B).

**Figure 3.**
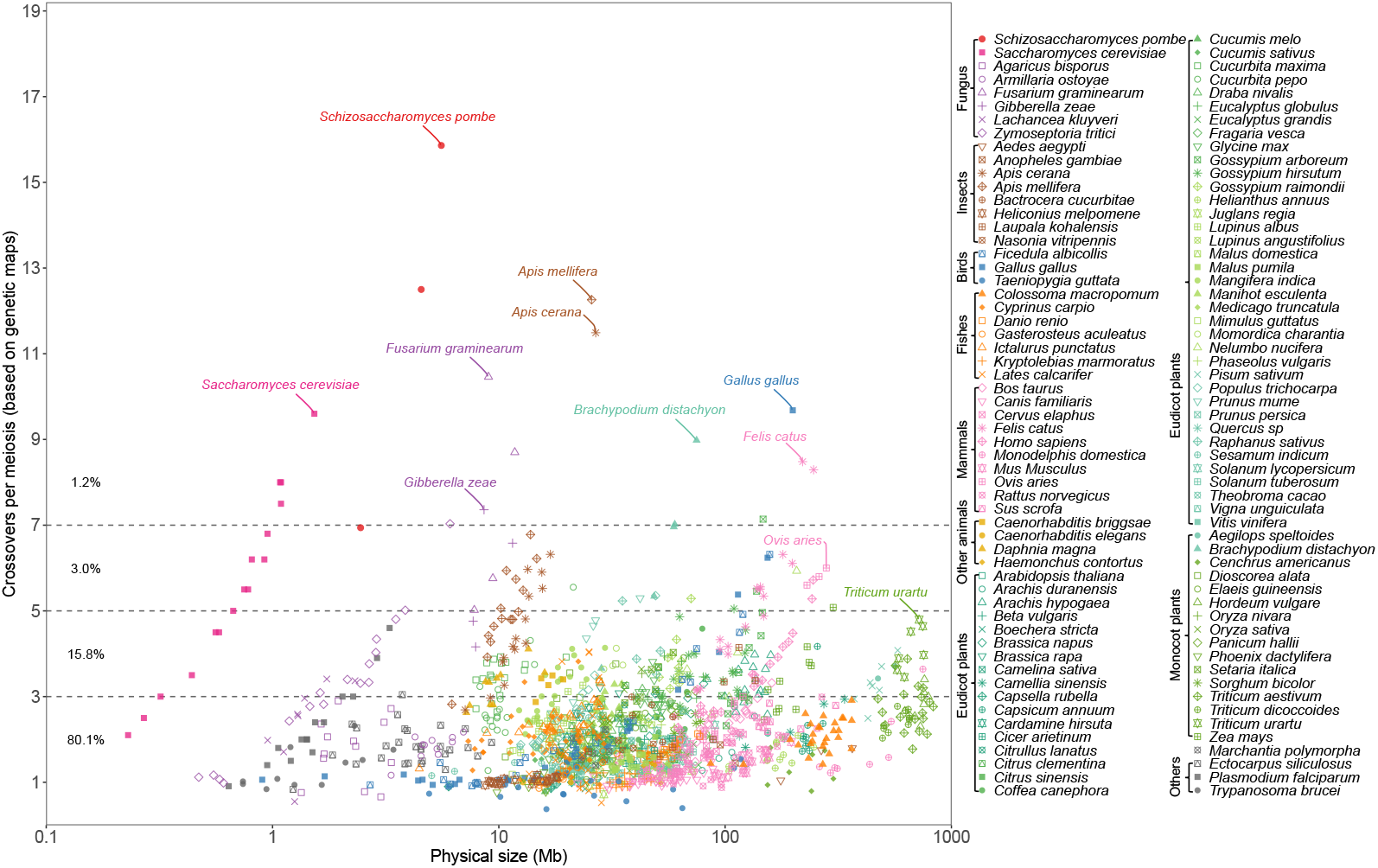
*S. pombe* has an exceptional high number of meiotic CO compared to other eukaryotes. The crossover number per meiosis (y-axis, based on genetic maps. 50 cM = 1 CO) of 114 species (8 fungi, 8 insects, 3 birds, 7 fishes, 10 mammals, 4 other animals, 54 eudicot plants, 16 monocot plants, 1 other plant and 3 other species) is plotted against the physical size (x-axis, DNA Mb, log scale) of each chromosome. The estimation of CO number of *S. pombe* (this study) is indicated by red dots. Plots separating the different eukaryotic clades are provided in Figure S7-8 and the source data in Table S6. An earlier version of this figure, with fewer species represented, was published by Fernandes et al. [8].

We then compared the crossover number per chromosome in a large range of eukaryotes (Figure 3). The number of crossovers per meiosis appears to be constrained: while at least one is always present, likely associated with the crucial role of COs in promoting balanced chromosome segregation at meiosis, larger numbers are rarely seen. In our survey of 1,547 chromosomes from 114 species, 80.1% of chromosomes have an average of less than three COs and 95.9% less than five (Figure 3, Table S6). CO numbers tends to be correlated with the physical size of the chromosomes within a species (Figure S7-8), and like in *S. pombe*, are linearly correlated with a constant CO frequency per megabase in species such as *S. cerevisiae*, *Arabidopsis thaliana*, *Ovis aries*, *Gallus gallus*, and *Homo sapiens* (Figure S7-8). In sharp contrast, this is not the case when comparing chromosomes from different species: for example, the physical size of chromosome 3B of wheat (804 Mb, *Triticum aestivum*) is 2,977-fold longer than the chromosome 6 of budding yeast (0.27 Mb, *Saccharomyces cerevisiae*), although they have similar genetic sizes (2.8 vs. 2.5 COs per meiosis, Figure 3). Several species have chromosomes with unusually large genetic sizes, like chromosome 1 of European honeybee (12.3 COs, *Apis mellifera*, Insects), chromosome 2 of *Fusarium graminearum* (10.5 COs, Fungus), and chromosome 1 of chicken (9.7 COs, *Gallus gallus*, Birds). *S. pombe* appears exceptional, with chromosomes I and II experiencing the highest number of crossovers and chromosome I having more than two times more crossovers than 99% of the eukaryotic chromosomes described so far. Chromosome III, the chromosome of *S. pombe* that undergoes the least recombination, is still involved in more COs than 98% of thus far studied chromosomes (Figure 3).

We next analyzed the CO distribution along *S. pombe* chromosomes. We observed that COs are relatively homogeneously distributed along chromosomes, largely distributing within the 2-fold standard deviation of the random distribution, without very hot or cold regions (Figure 2F, Figure S9). Notably, we do not detect a strong effect of the centromere on CO rates. However, it should be noted that the limited resolution of our study may hide strong local variation in recombination rates, including a probable suppression next to the centromere as observed in many other eukaryotes [16, 20–26].

In addition to COs, we identified 66 NCOs characterized by a 3:1 segregation profile of a single SNP, which corresponds to an average of 0.7 per tetrad (Figure 2C, 2E, and Figure S3). In total, we analyzed the segregation of 13,975 SNPs in 90 tetrads, which resulted in an estimation of an average conversion rate of 0.47% per base pair per meiosis. With a genome size of 12.57 Mb, this predicts than on average 59 kb are converted per meiosis. In comparison, in *S. cerevisiae* 66 NCOs with a median of 1.8 kb were observed, corresponding to 119 kb undergoing gene conversion per meiosis. This ca. two-fold difference suggests that either the number of NCOs is lower in *S. pombe* than *S. cerevisiae* and/or that the NCO-associated gene conversion tract length is shorter in *S. pombe* compared to *S. cerevisiae*. If we assume that the NCO-associated gene conversion tract length is similar in both species, this would predict an average number of 33 NCOs per meiosis, very close to our estimation of CO number (35 per meiosis), suggesting that DSBs are repaired as either COs and NCOs with similar frequencies in *S. pombe*.

We also observed 37 gene conversions (GCs) with a 4:0 allelic segregation. Remarkably, 13 and 8 of these GCs occurred in only two independent tetrads of the pombe2 population, and a common GC occurred in 16 tetrads of the pombe4 population (Figure 2C, Figure S3 and S10). Although these events can result from the GC-mediated repair of two double-strand breaks at the same locus on two sister chromatids during meiotic prophase, such a scenario is unlikely. Instead, these events more likely reflect mitotic gene conversion events that preceded meiotic entry [16–18]. However, the diploid phase of *S. pombe* is limited since haploid cells mate and sporulate on the same culture medium. The fact that two tetrads out of 90 contain 21/37 4:0 segregating NCOs while all the other tetrads have never more than three such NCOs could reflect an elevated instability at the diploid stage potentially promoted by an uncontrolled *Spo11* expression prior to the pre-meiotic S phase in a small fraction of cells from the population. The recurrent 4:0 segregating NCOs in the 16 tetrads of the pombe4 population could reflect either an elevated meiotic or mitotic instability of a given locus.

Along chromosomes, COs are not independently distributed in most eukaryotes, as CO interference prevents close positioning [27, 28]. Based on tetrad analysis, *S*. *pombe* has been shown to lack CO interference [9]. To further explore CO interference in *S*. *pombe*, we first examined the distribution of the distance between adjacent COs on the same chromosome, comparing with the expected distribution in the absence of CO interference (Figure 4A–B, Figure S11A–F). We simulated the expected distribution by using the same marker list, and kept the same sample size and CO number for each chromosome, tetrad and population. The observed distribution was found to be remarkably similar to that expected for independent COs, with numerous pairs of COs placed less than 100 kb apart. This contrasts with observations in *S. cerevisiae* in which close COs are virtually absent [16] (Figure 4A-B).

**Figure 4.**
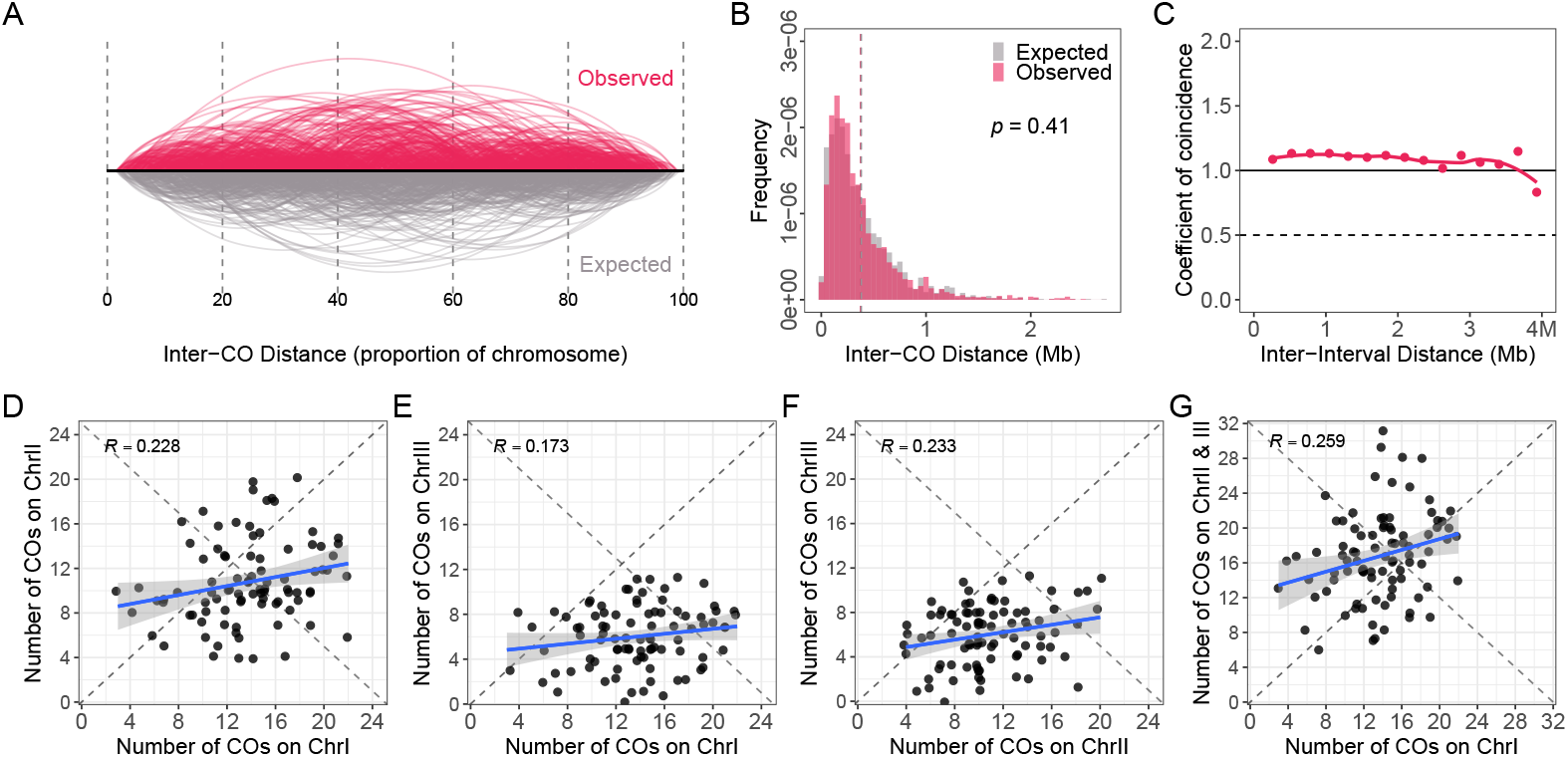
Absence of CO interference but presence of co-variation of CO frequencies in *S. pombe*. (A) Comparison of the observed (red) and expected (grey) distribution of inter-CO distances along the proportional scale of chromosomes. (B) Distribution of observed (red) and expected in absence of interference (grey) distances between COs. The expected CO list was obtained as one simulation by sampling the observed CO number per chromosome and tetrad in the corresponding scale of genomic regions covered by EMS-induced SNPs. The p-value from a Mann-Whitney test between the expected and observed distributions is shown. (C) The CoC curves of the observed COs. Chromosomes were divided into 16 intervals, and the coefficient of coincidence was calculated for each possible pair of intervals (observed frequency of concomitant CO in both intervals/product of frequency of CO in each interval. A coefficient of coincidence of 1 indicates absence of interference). The coefficient of coincidence was averaged for pairs of intervals separated by the same distance (e.g., pairs of adjacent intervals), and plotted versus the distance between the center of the two intervals. (D–G) Correlation analysis of the numbers of COs on different chromosomes in the same tetrad. The blue line indicates the linear regression, and the grey zone represents the 95% confidence interval.

To further describe CO interference, we performed coefficient of coincidence (CoC) analysis, which is the ratio of observed frequency of double-COs to the frequency of expected double-COs in a pair of defined intervals [29, 30]. We found that the CoC curve is flat (close to 1), supporting a complete absence of CO interference in *S. pombe* (Figure 4C, Figure S11G–I). It should be noted that because of limited resolution, we cannot exclude CO interference acting at short distances [31]. Intriguingly, the CoC value is slightly above 1 all along the chromosomes, suggesting that the presence of a CO favors the occurrence of another CO in the same chromosome. We then tested the existence of covariation of CO numbers between chromosomes within nuclei, as observed in humans and other eukaryotes [32]. We observed that CO number tends to covary across chromosomes within individual tetrads, (Figure 4D–G, Figure S11J–M). The numbers of COs per chromosome are positively correlated between chromosomes (Figure 4D–F) and also between wo groups of chromosomes with comparable physical lengths (ChrI v.s. ChrII and III, Figure 4G). This indicates that if one cell has a large number of COs on one chromosome, it tends to have also a large number of COs on the other chromosomes. This phenomenon could account for the CoC value being above 1 and suggests that a limiting factor for CO formation stochastically varies between cells, causing some cells to have more COs than others, either conjointly on different chromosomes or at different positions on the same chromosome. As the SC and interference are absent, we speculate that variation in DSB frequencies may be the source of cell-to-cell variations in COs, potentially through variation in the condensation of the loop-axis structure [32].

Finally, we explored chromatid interference. As described above, when considering two adjacent COs, three types of outcomes would be expected depending on how many chromatids are affected (2C-DCOs, 3C-DCOs and 4C-DCOs, Figure S6). In the absence of chromatid interference, DCOs are randomly distributed among the four chromatids of the bivalent, and the three types of DCOs will give a ratio of 1:2:1 (2C:3C:4C DCOs) [33]. If the chromatid implicated in a CO affects somehow the choice of the chromatid of the adjacent CO, we would expect a deviation from this ratio. In our *S. pombe* tetrad analysis, the ratio of 2C:3C:4C DCOs for each individual chromosome (1,012, 768 and 399 pair of COs were analyzed separately) and the whole genome (2,179 CO pairs), did not significantly deviate from 1:2:1 (Chi-Square test, Table 1). To explore the potential centromere effect on chromatid interference, we further quantified the deviation from the expect 1:2:1 ratio of 2C:3C:4C DCOs for two groups of DCOs on the same chromosome arms or that span the centromere, separately. In all cases, no significant deviation was detected (Table 1). These results showed an absence of chromatid interference in fission yeast, confirming the previous report which had less statistical power [9].

**Table 1.** Chromatid interference is absent in *S. pombe*.

|  | Whole chromosome |  | Same arm |  | Different arm |  |
| --- | --- | --- | --- | --- | --- | --- |
|  | Total DCOs | Observed 2C:3C:4C ratio | Total DCOs | Observed 2C:3C:4C ratio | Total DCOs | Observed 2C:3C:4C ratio |
| I | 1012 | 247:502:263<br>($p = 0.821$) | 930 | 222:467:241<br>($p = 0.821$) | 82 | 25:35:22<br>($p = 0.668$) |
| II | 768 | 184:384:200<br>($p = 0.821$) | 688 | 164:346:178<br>($p = 0.821$) | 80 | 20:38:22<br>($p = 0.861$) |
| III | 399 | 95:187:117<br>($p = 0.668$) | 338 | 87:156:95<br>($p = 0.668$) | 61 | 8:31:22<br>($p = 0.479$) |
| Total | 2179 | 526:1073:580<br>($p = 0.668$) | 1956 | 473:969:514<br>($p = 0.668$) | 223 | 53:104:66<br>($p = 0.668$) |

\* 2C DCOs mean two COs use the same non-sister chromatids; 3C DCOs mean two COs share one chromatid; 4C DCOs mean two COs use the different pair of non-sister chromatids.
\* Chi-Square test of goodness-of-fit was performed for each ratio, and none significant deviation from the 1:2:1 ratio was detected. The significant p value was corrected by the fdr method.

In summary, regulation of meiotic recombination appears to be relaxed in *S. pombe*, resulting in an exceptionally high number of crossovers per chromosome, a random distribution of crossover among chromosomes and absence of crossover interference. The high frequency of COs, even though randomly distributed, makes it highly unlikely that a given chromosome will lack a CO, thus ensuring balanced chromosome distribution. It is intriguing that this simple strategy to ensure the obligate crossover was retained in very few species, while most eukaryotes have a low number of COs and evolved sophisticated mechanisms to ensure their non-random distribution among and along chromosomes, notably to ensure the obligate COs. This suggests that a strong evolutionary constraint imposes a low CO number in eukaryotes and that *S. pombe* either escaped this constraint or is in an evolutionary dead-end.

## Materials and Methods

### Strain construction, mating, growth conditions and tetrad dissection

Media and methods for studying and mutagenizing *S. pombe* were as described in [34]. *S. pombe* strains PR109 (h-leu1-32 ura4-D18) and PR110 (h+ leu1-32 ura4-D18) were used for three successive rounds of mutagenesis with ethylmethane sulfonate mutagenesis (EMS from Sigma) based on [35]. Briefly, *S. pombe* cells were inoculated in liquid YES medium overnight at 32 °C, diluted to 2-5 x 10exp6 cells/ml the next morning and grown for 6 hours. Next, cells were resuspended in liquid EMM at 1 x 10exp8 cells/ml and 1 ml was treated or not with 20 microliters of EMS (i.e., 2% final) for 3 h in 10 ml tubes at 32 °C. Next, cells were washed three times with 5 ml of EMM, resuspended in 1 ml of EMM, plated on YES plates and incubated at 32 °C until colonies formed. After such EMS treatment, cell viability was around 20%. Five independent clones of the first round of mutagenesis were at the root of two subsequent similar rounds of mutagenesis. Each clone used was checked for its ability to mate and sporulate. Eventually, five mutagenized clones from each of the PR109 and PR110 backgrounds were sequenced to identify *de novo* mutations and determine the optimal combinations of mutation patterns for recombination analyses. The following h-x h+ crosses were selected and the corresponding tetrads dissected: BLP49 (A) x BLP23 (I), 29 tetrads; BLP59 (C) x BLP19 (H), 30 tetrads and BLP64 (D) x BLP33 (K), 33 tetrads.

### Genomic DNA extraction and sequencing

*S. pombe* spores were grown to saturation in 1.6 ml of YES medium at 32 °C in 96-well plates (2 ml deep well plates). Cells were harvested by centrifugation, washed once with TE and lyzed during two hours at 37 °C using 30 units of zymolyase in Y1 Qiagen buffer. Samples were centrifuged at 2,700 g for five minutes and supernatants were treated with 0.5 mg/ml of RNaseA for 10 minutes at room temperature. DNA was next purified from these lysates using the E-Z 96™ Tissue DNA Kit from Omega Bio-tek following the manufacturer’s instructions except for DNA binding to the columns that was done by gravity flow instead of centrifugation. The sequencing library was then prepared for 2×150 bp HiSeq 3000 Illumina sequencing [36]. The Mendelian segregation of the mating type locus in each tetrad was verified by PCR as in [37].

### High quality EMS-induced mutation marker calling

The reference genome and genomic annotations of the fission yeast *Schizosaccharomyces pombe* were downloaded from the PomBase database (https://www.pombase.org) [38, 39]. The whole genome resequencing short reads of parental samples of the three independent populations (Pombe2_Tet, Pombe4_Tet and Pombe9_Tet, Table S2) were aligned to the reference genome using BWA v0.7.15-r1140 [40] with the default parameters. A command set of Sambamba v0.6.8 [41] was used to format the read alignment file. The variant calling and filtering processes were largely performed following a previously described pipeline [26, 42–45]. First, variations were called for each individual parental sample by inGAP-family [44]. Then, non-allelic, low-quality and artificial variants were selected and filtered by checking the effects of tandem repeats, small indels and structural variations predicted by Tandem Repeats Finder v4.09 [46] and inGAP-family, separately. Finally, mutations (G to A and C to T) specific to each parental genome were kept as high-quality markers (Table S3) for subsequent analysis.

### Identification of meiotic recombination events

The sequencing reads of the 92 tetrads (Table S1) were aligned to the reference genome by BWA and then formatted by Sambamba as described before. The sequencing depths were measured using mosdepth v0.3.3 [47] with “-n --fast-mode --by 100”. For each spore, high-quality mutation markers were genotyped by inGAP-family. The sequencing depths, reads mapping ratio, the covered marker number and the allele frequency of mutation markers were estimated and examined to remove potentially contaminated and problematic samples (Table S1). Two tetrads (Pombe9_Tet_10 and Pombe9_Tet_14) were discarded, because more than one problematic spore was detected (Table S1). During the genotyping process, markers that were not covered or poorly covered in only one of the four spores was imputed and re-genotyped based on the segregation pattern. Meiotic CO events were defined as two nearby markers showing different genotype patterns in the four spores and with a segregation ratio of 2:2. GC events were identified as markers with 3:1 or 4:0 segregation pattern among the four spores. For each tetrad, identified CO and GC events were manually checked by visualization of the marker genotype and chromosomal genotypes. For those unsolved double-CO events (Figure S5), which are two COs occurred within the interval of two adjacent markers and involve all four chromatids, and the pair of homolog chromosomes that CO occurred cannot be differentiated, these unsolved DCOs were removed in the subsequent CO and chromatid interference analysis.

Sparse coverage and unreliable genotyping may cause biased calling of COs. Therefore, we examined the effect of sequencing depths on CO detection. We found that the resolution and numbers of detected COs were independent of sequencing depth (mean = 19.1x, Pearson’s correlation coefficient, *R* = 0.03 and −0.03), suggesting sufficient sequencing depths and the absence of bias in CO identification among tetrads (Figure S3B and S12).

### CO and chromatid interference analysis

To simulate the CO landscape in the absence of interference, COs were randomly sampled in the genomic range of first and last markers in the population along chromosomes. To match the contribution of COs per chromosome and tetrad, we kept the exact same sample size in the simulation. The distances between simulated COs were calculated as the observed dataset. The CoC analysis was performed using MADpattern v.1.1 [29, 30]. For the chromatid interference analysis, the Chi-Square test of goodness-of-fit was performed to evaluate the deviation from the random 1:2:1 ratio for each observed 2C:3C:4C ratio, using the chisq.test function (stats v3.6.2 package) in R environment. Adjacent double COs were not included in the CO and chromatid interference analysis.

## Data availability

The raw sequencing data can be accessed in ArrayExpress under the accession numbers E-MTAB-12514 and E-MTAB-12516. The list of identified COs and GCs can be found in Table S4.

## Acknowledgements

We would like to thank the Max Planck Genome centre for library preparation and sequencing. We thank Neysan Donnelly for proofreading the manuscript. This work was supported by core funding from the Max Planck Society and an Alexander von Humboldt Fellowship to Q.L. B.L. lab was supported by the ANR grant ANR-18-CE12-0013-01.

## Author contributions

Q.L., B.L. and R.M. designed the research. B.L., L.M. and M.G. generated the biological material. Q.L., B.L. and R.M. analyzed the data. Q.L. and R.M. wrote the article with input from B.L.

## Supplemental Figure legends

**Figure S1. Chromosomal distribution of EMS-induced SNPs.**

(A) The distribution of EMS-induced SNPs along chromosomes. Comparison of the expected and observed distribution of inter-marker distances in the whole population (B), Pombe2_Tet (C), Pombe4_Tet (D) and Pombe9_Tet populations (E), separately.

**Figure S2. Interval length distribution of COs in all chromosomes together and in each replicate population.**

The distribution of CO interval lengths in the whole population (A), Pombe2_Tet (B), Pombe4_Tet (C) and Pombe9_Tet (C) populations, separately. The black dashed line indicates the median value. The median value is shown in the corresponding plot.

**Figure S3. Distribution of CO and GC numbers and sequencing depths in all chromosomes together, each chromosome individually, and in the replicate population.**

(A) Comparison of the CO and GC numbers in each replicate population. (B) Comparison of the CO and GC numbers and sequencing depths in each replicate population. (C) The distribution of CO numbers in each chromosome and replicate population. (D) The distribution of GC numbers in each chromosome and replicate population.

**Figure S4. Comparison of CO numbers per chromosome between fission and budding yeast.** Correlation analysis of CO numbers and chromosome lengths (Mb of DNA). The mean number of COs is shown with the standard deviation. The dots were connected by dashed lines with corresponding colors. The fitted linear regression is indicated by the grey line.

**Figure S5. The genotype profile of a tetrad with a double CO in a single interval.**

The chromosomal distribution of genotype of mutation markers in each of the four spores. The detected COs and GCs are indicated by dashed lines and triangles separately. The chromosomes are color-coded according to the parental genotypes. The red arrows mark the unsolved DCO.

**Figure S6. Schematic representation of the four types of double-crossover events.**

The double-crossover events are classified based on the number of involved chromatids. The black dashed lines indicate two adjacent EMS-induced SNPs. When double-crossover events occur in the interval of two adjacent EMS-induced SNPs, 2C-DCO is undetectable, 3C-DCO would be detected as a single CO, and 4C-DCO would be not affected.

**Figure S7. The comparison of crossovers per meiosis with physical size of chromosomes in each group of eukaryotes.**

The crossover number per meiosis (y-axis, based on genetic map, linear scale) of 114 species (8 fungi, 8 insects, 3 birds, 7 fishes, 10 mammals, 4 other animals, 54 eudicot plants, 16 monocot plants, 1 other plant and 3 other species) is plotted against the physical size (x-axis, Mb) of each chromosome. The dot was connected by dashed lines with corresponding colors. The fitted linear regression is indicated by the grey line.

**Figure S8. The comparison of crossovers per meiosis with physical size of chromosomes in four groups of eukaryotes.**

The crossover number per meiosis (y-axis, based on on genetic map, linear scale) of 114 species (6 fishes, 9 mammals, 53 eudicot plants and 16 monocot plants, with chromosome lengths smaller than 80, 350, 250 and 100 Mb, respectively) is plotted against the physical size (x-axis, Mb) of each chromosome. The dot was connected by dashed lines with corresponding colors. The fitted linear regression is indicated by the grey line.

**Figure S9. The chromosomal distribution of detected and expected COs.**

The detected (pink and red) and expected (grey) CO distribution along chromosomes with 100 kb (A) and 200 kb (B) sliding window size and 10 kb step size. The expected CO list was obtained as 200 rounds of simulation by sampling the observed CO number per chromosome and tetrad in the corresponding scale of genomic regions covered by EMS-induced SNPs. The expected CO distribution is shown with 2-fold standard deviation (grey shading). The centromeric regions are indicated by grey shading.

**Figure S10. Genotype profile of two tetrads with extreme number of GCs.**

The chromosomal distribution of the genotypes of mutation markers in each of the four spores. The detected COs and GCs are indicated by dashed lines and triangles, separately. The chromosomes are color-coded according to the parental genotypes.

**Figure S11. CO interference and covariation analysis in each replicate population.**

(A–C) Comparison of the observed and expected (grey) chromosomal distributions of inter-CO distances in each replicate population. (D–F) Comparison of the observed and expected (grey) inter-CO distances in each replicate population. The expected CO list was obtained as one simulation by sampling the observed CO number per chromosome and tetrad in the corresponding scale of genomic regions covered by EMS-induced SNPs. (G–I) The CoC curves of observed COs in each replicate population. (J–M) Comparison of the numbers of COs between chromosomes in individual tetrads and each replicate population.

**Figure S12. Comparison of distributions of CO interval lengths and sequencing depths in each replicate population.**

The median values are indicated by dashed lines.

**Table S1. Summary of the sequencing data of the progeny populations.**

**Table S2. Summary of the sequencing data of the parental strains.**

**Table S3. The table of detected mutations among the parental strains.**

**Table S4. The list of detected COs and GCs.**

**Table S5. Estimation of genetic lengths of chromosomes.**

**Table S6. Genetic sizes in eukaryotes**

