## Supplementary material for "Remarkably high rate of meiotic recombination in the fission yeast *Schizosaccharomyces pombe*": Figure S1 to S12

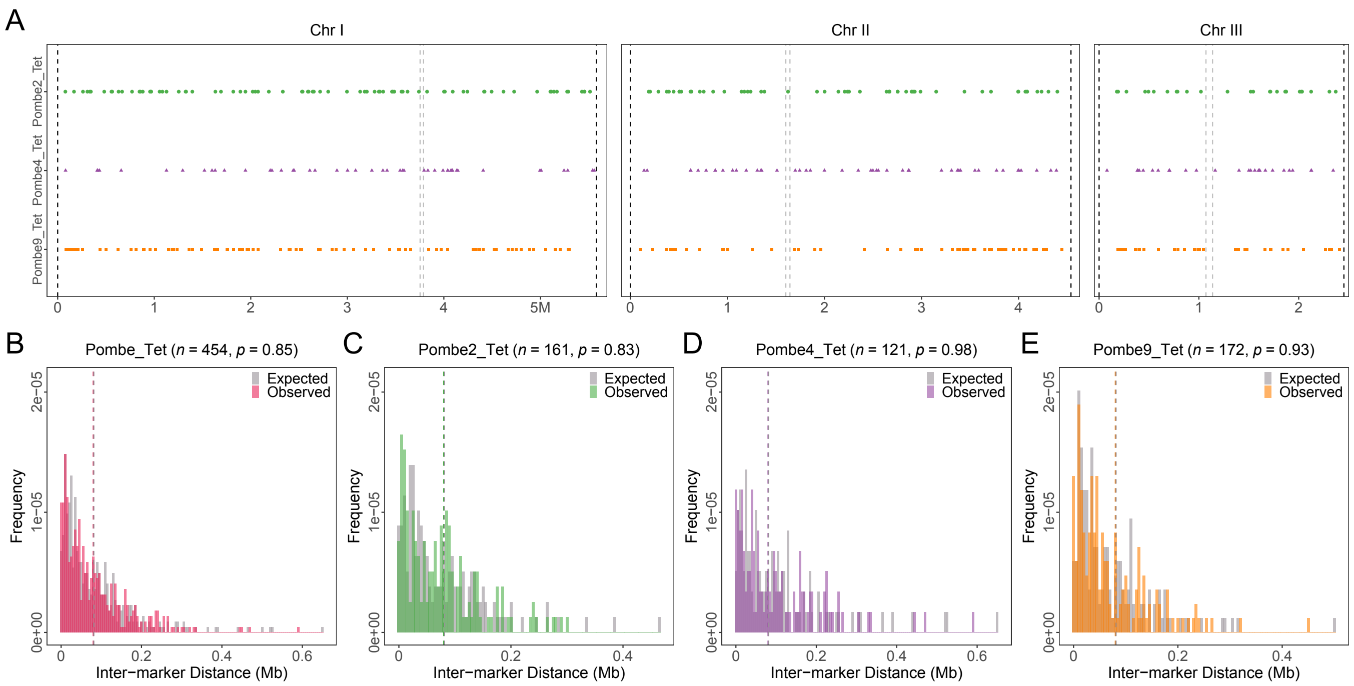


**Figure S1. Chromosomal distribution of EMS-induced SNPs.**

(A) The distribution of EMS-induced SNPs along chromosomes. Comparison of the expected and observed distribution of inter-marker distances in the whole population (B), Pombe2_Tet (C), Pombe4_Tet (D) and Pombe9_Tet populations (E), separately.

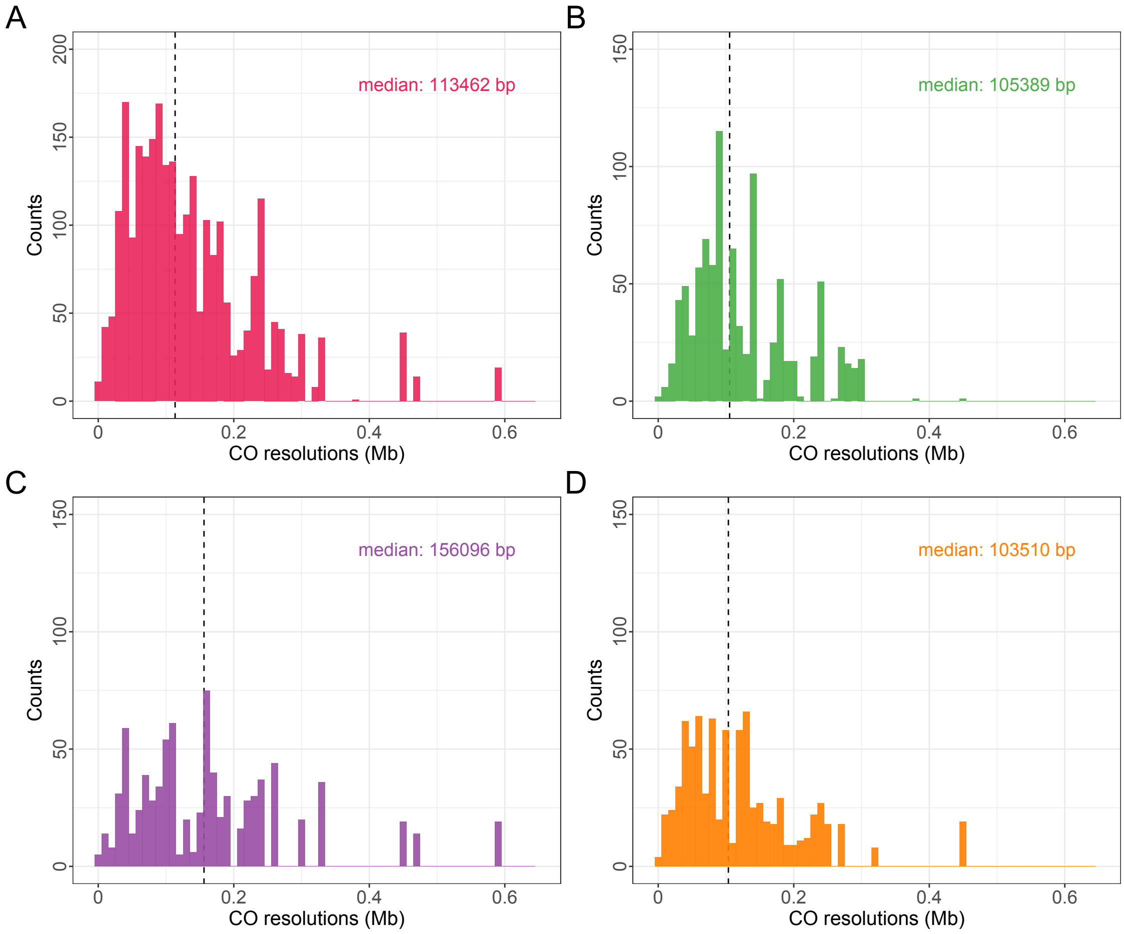


**Figure S2. Interval length distribution of COs in all chromosomes together and in each replicate population.**
The distribution of CO interval lengths in the whole population (A), Pombe2_Tet (B), Pombe4_Tet (C) and Pombe9_Tet (C) populations, separately. The black dashed line indicates the median value. The median value is shown in the corresponding plot.

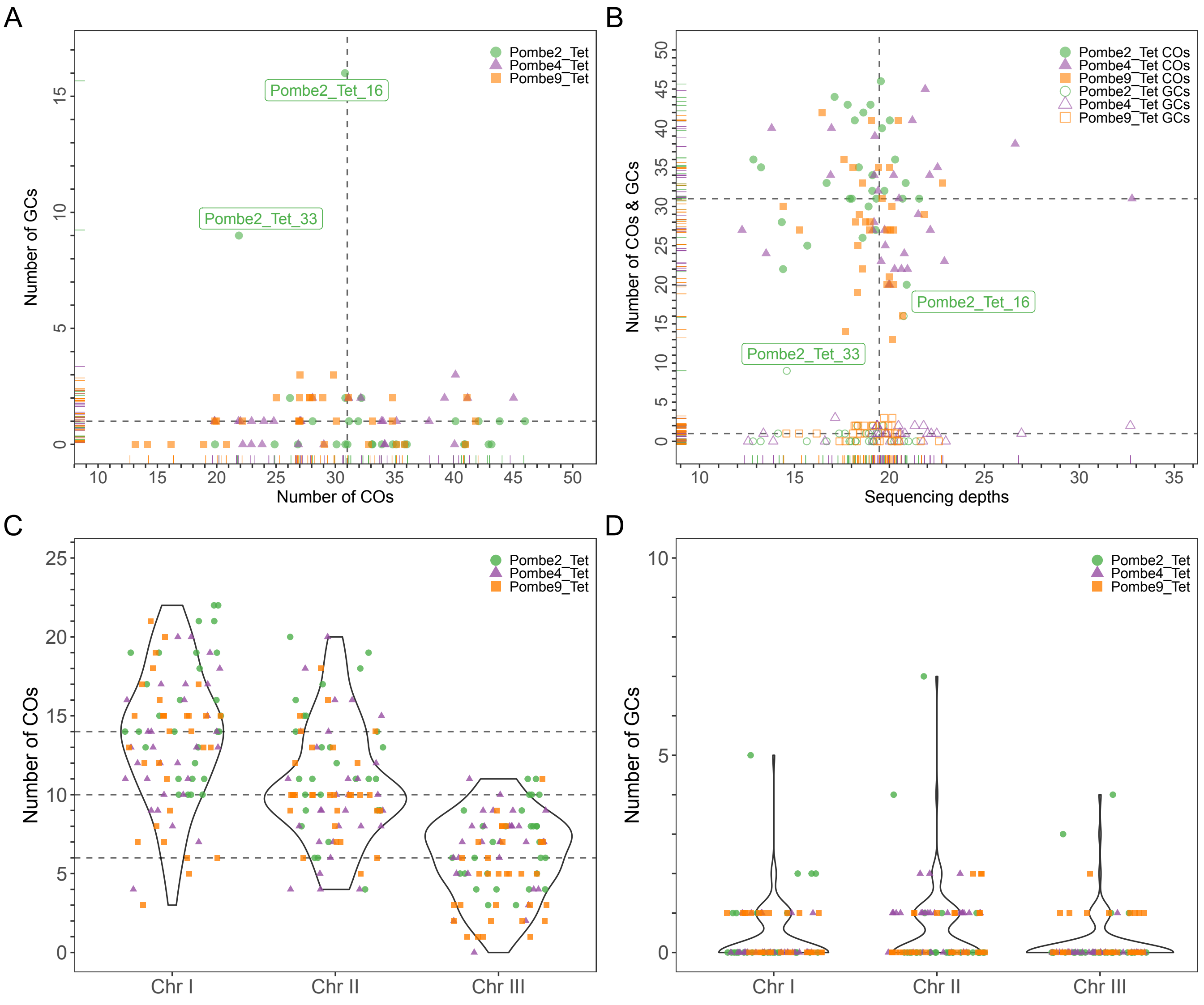


**Figure S3. Distribution of CO and GC numbers and sequencing depths in all chromosomes together, each chromosome individually, and in the replicate population.**

(A) Comparison of the CO and GC numbers in each replicate population. (B) Comparison of the CO and GC numbers and sequencing depths in each replicate population. (C) The distribution of CO numbers in each chromosome and replicate population. (D) The distribution of GC numbers in each chromosome and replicate population.

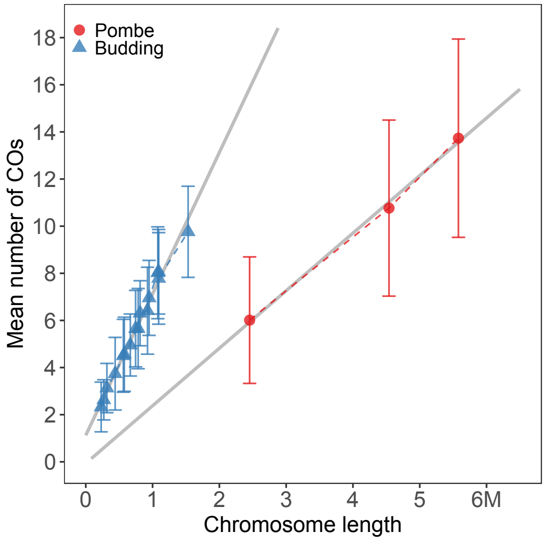


**Figure S4. Comparison of CO numbers per chromosome between fission and budding yeast.**
Correlation analysis of CO numbers and chromosome lengths (Mb of DNA). The mean number of COs is shown with the standard deviation. The dots were connected by dashed lines with corresponding colors. The fitted linear regression is indicated by the grey line.

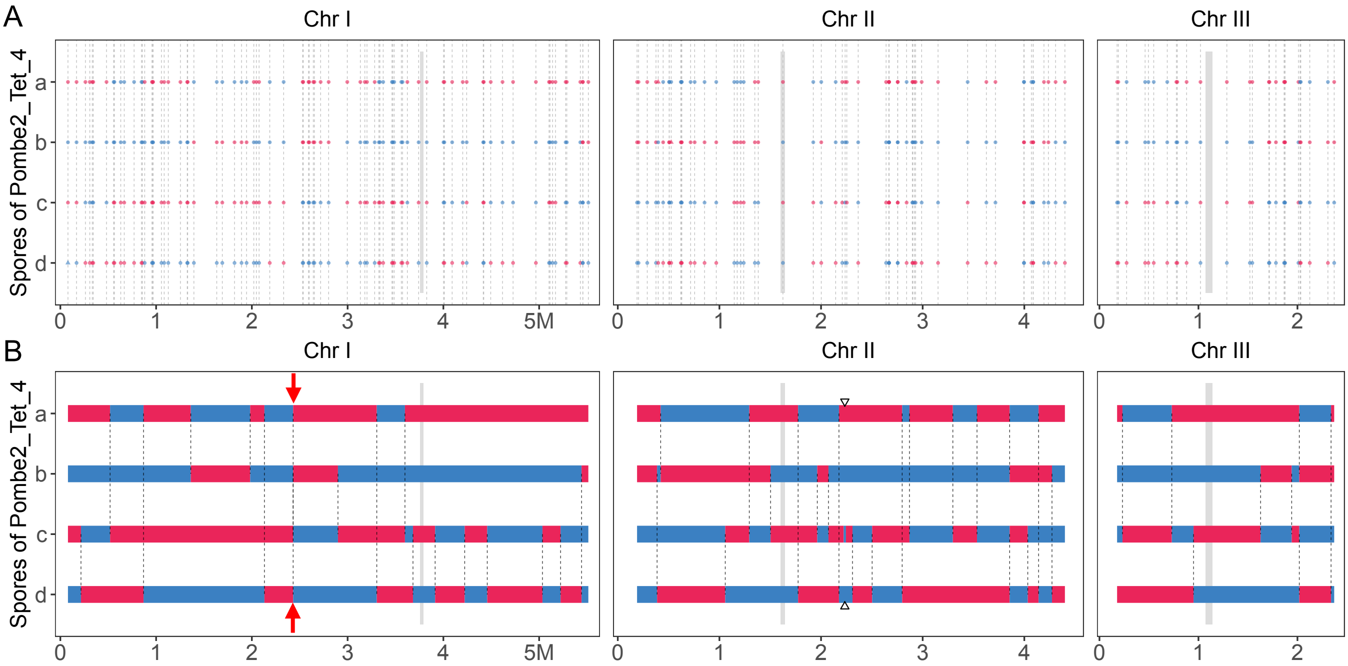


**Figure S5. The genotype profile of a tetrad with a double CO in a single interval.**
The chromosomal distribution of genotype of mutation markers in each of the four spores. The detected COs and GCs are indicated by dashed lines and triangles separately. The chromosomes are color-coded according to the parental genotypes. The red arrows mark the unsolved DCO.

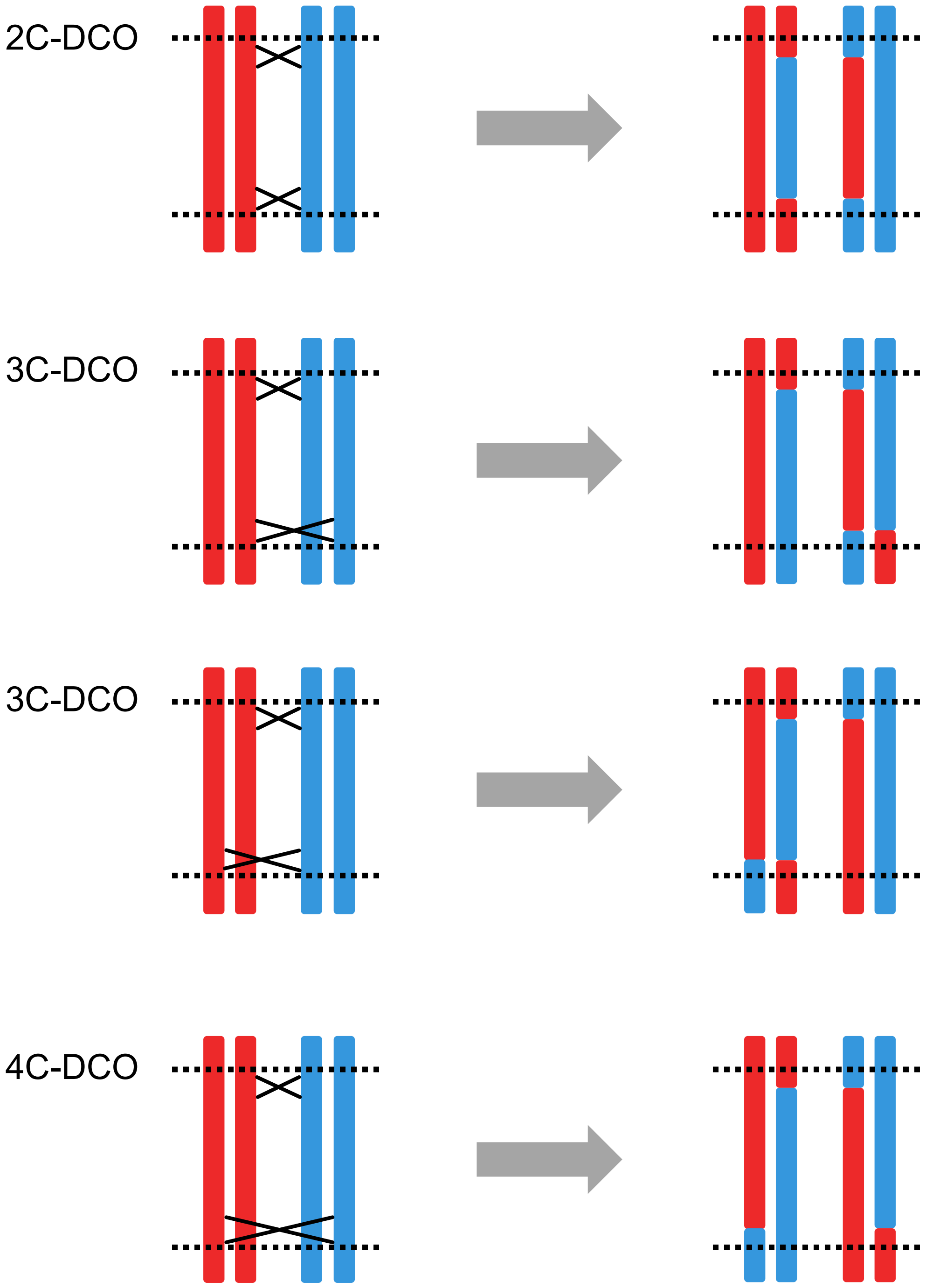


**Figure S6. Schematic representation of the four types of double-crossover events.** The double-crossover events are classified based on the number of involved chromatids. The black dashed lines indicate two adjacent EMS-induced SNPs. When double-crossover events occur in the interval of two adjacent EMS-induced SNPs, 2C-DCO is undetectable, 3C-DCO would be detected as a single CO, and 4C-DCO would be not affected.

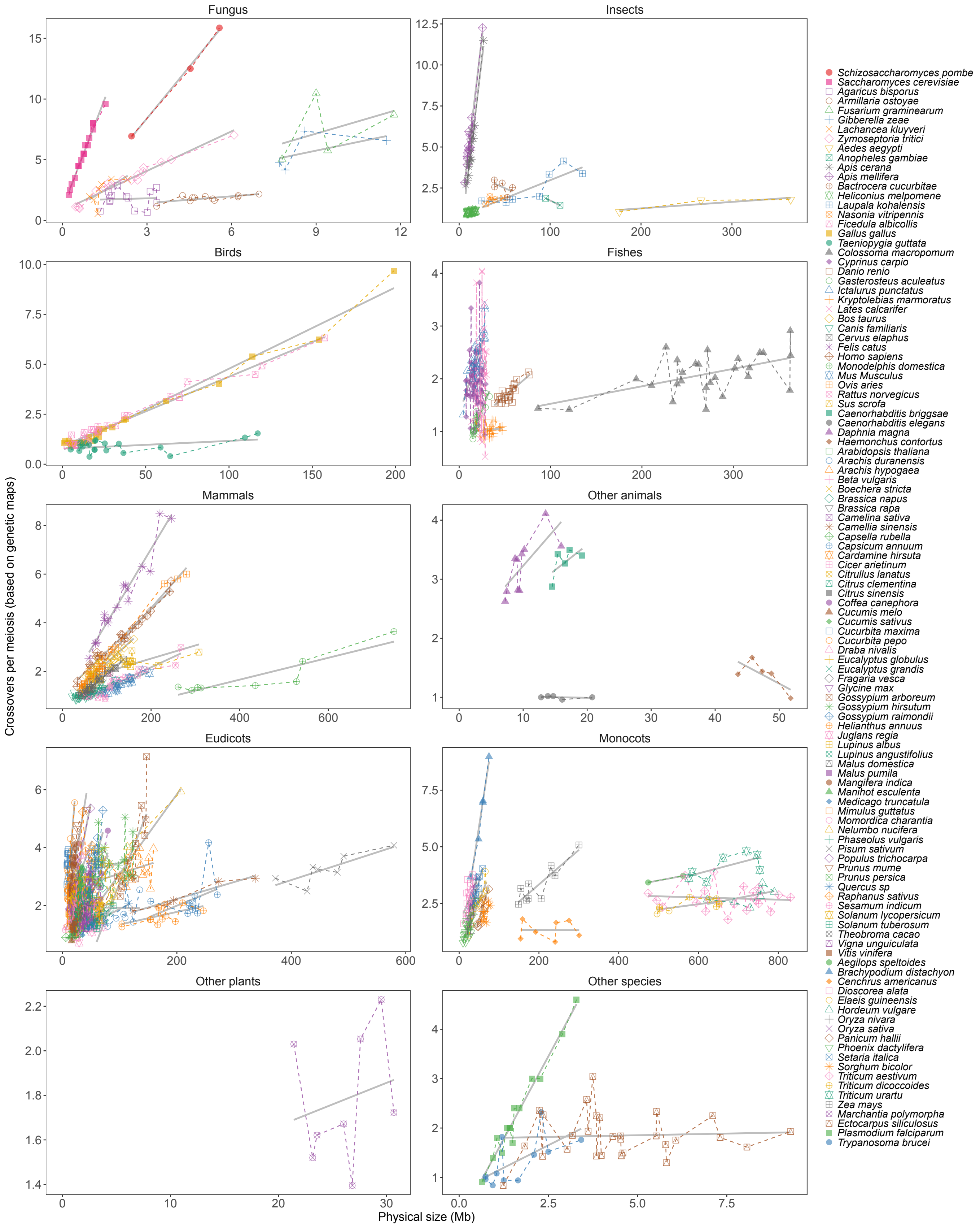


**Figure S7. The comparison of crossover per meiosis with physical size of chromosomes in each group of eukaryotes.**

The crossover number per meiosis (y-axis, based on genetic map, linear scale) of 114 species (8 fungi, 8 insects, 3 birds, 7 fishes, 10 mammals, 4 other animals, 54 eudicot plants, 16 monocot plants, 1 other plant and 3 other species) is plotted against the physical size (x-axis, Mb) of each chromosome. The dot was connected by dashed lines with corresponding colors. The fitted linear regression is indicated by the grey line.

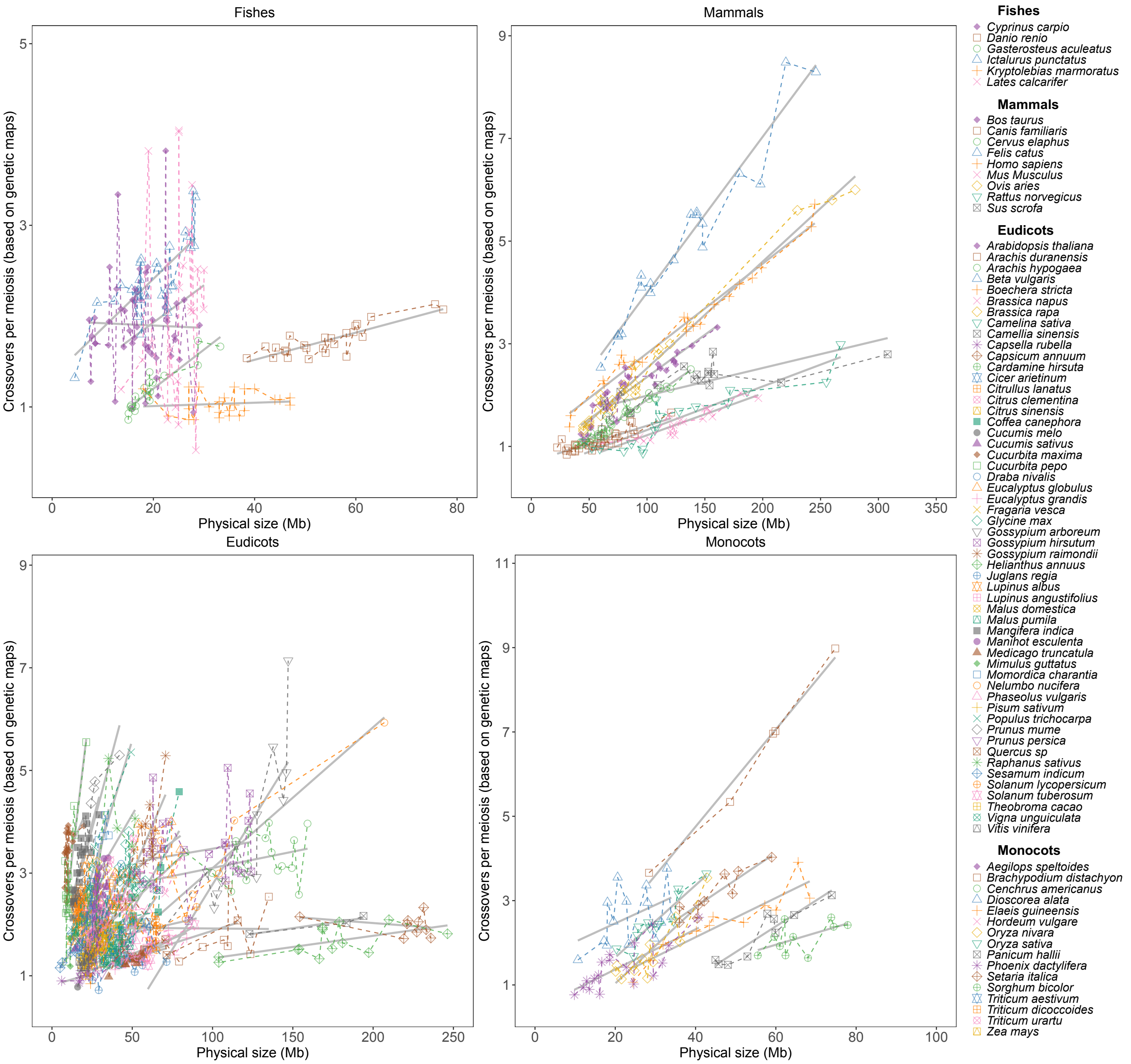


**Figure S8. The comparison of crossover per meiosis with physical size of chromosomes in four groups of eukaryotes.**

The crossover number per meiosis (y-axis, based on on genetic map, linear scale) of 114 species (6 fishes, 9 mammals, 53 eudicot plants and 16 monocot plants, with chromosome lengths smaller than 80, 350, 250 and 100 Mb, respectively) is plotted against the physical size (x-axis, Mb) of each chromosome. The dot was connected by dashed lines with corresponding colors. The fitted linear regression is indicated by the grey line.

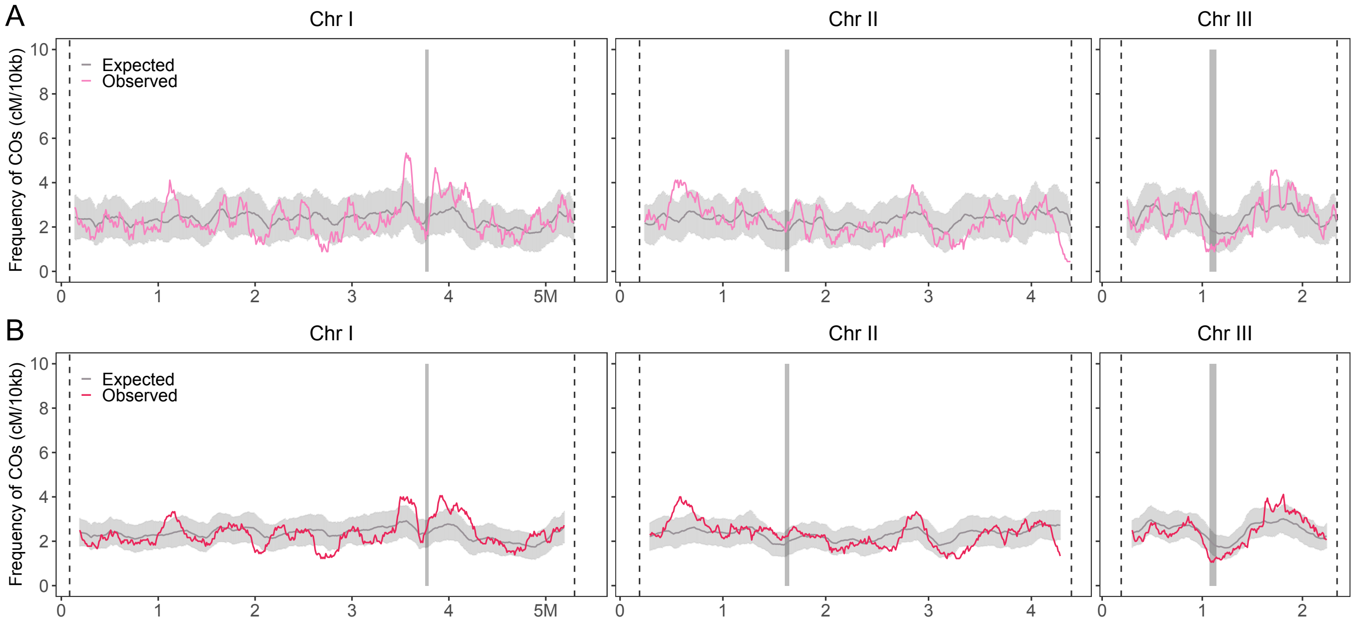


**Figure S9. The chromosomal distribution of detected and expected COs.**
The detected (pink and red) and expected (grey) CO distribution along chromosomes with 100 kb (A) and 200 kb (B) sliding window size and 10 kb step size. The expected CO list was obtained as 200 rounds of simulation by sampling the observed CO number per chromosome and tetrad in the corresponding scale of genomic regions covered by EMS-induced SNPs. The expected CO distribution is shown with 2-fold standard deviation (grey shading). The centromeric regions are indicated by grey shading.

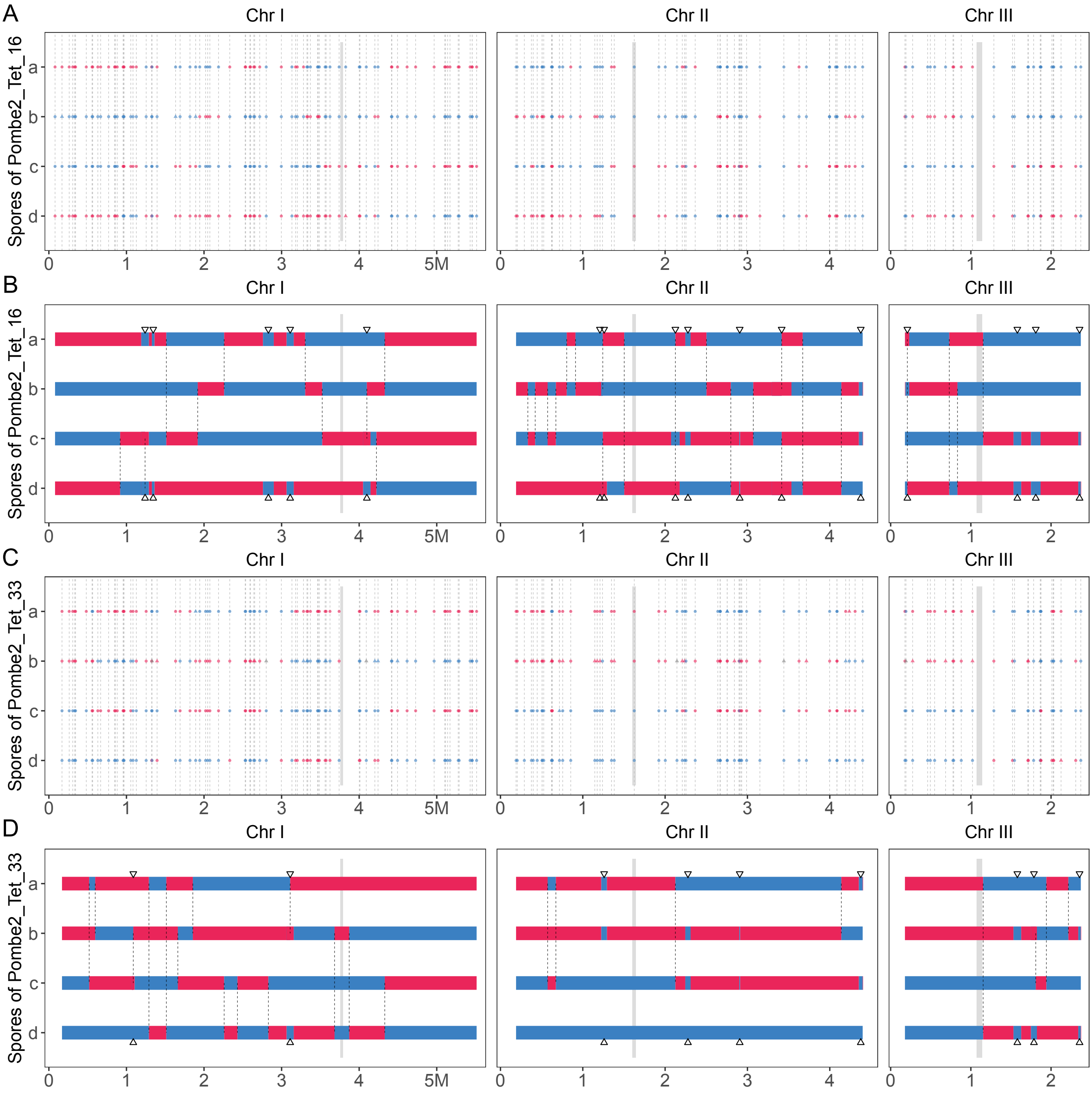


**Figure S10. Genotype profile of two tetrads with extreme number of GCs.**
The chromosomal distribution of the genotypes of mutation markers in each of the four spores. The detected COs and GCs are indicated by dashed lines and triangles, separately. The chromosomes are color-coded according to the parental genotypes.

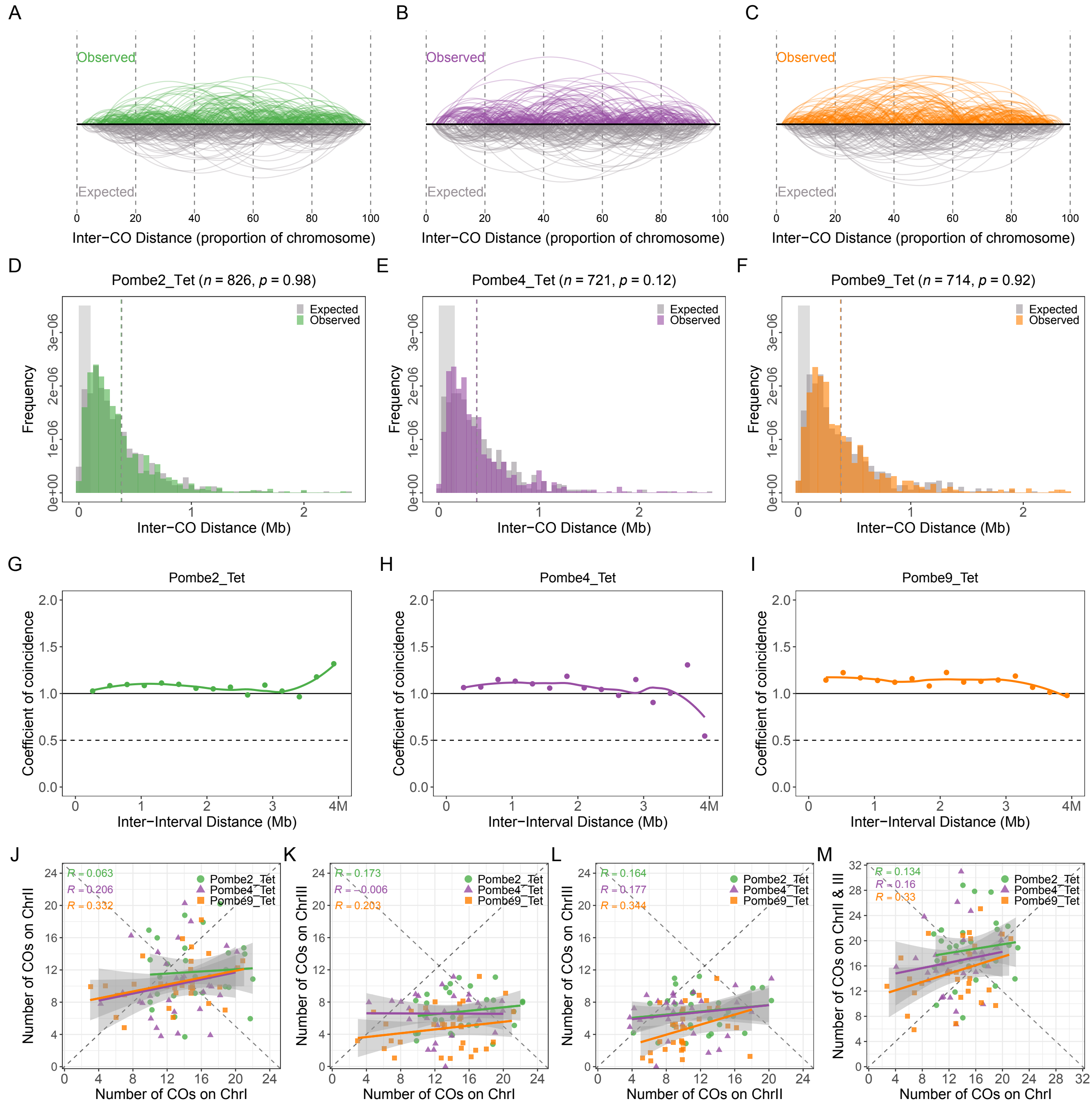


**Figure S11. CO interference and covariation analysis in each replicate population.** (A–C) Comparison of the observed and expected (grey) chromosomal distributions of inter-CO distances in each replicate population. (D–F) Comparison of the observed and expected (grey) inter-CO distances in each replicate population. The expected CO list was obtained as one simulation by sampling the observed CO number per chromosome and tetrad in the corresponding scale of genomic regions covered by EMS-induced SNPs. (G–I) The CoC curves of observed COs in each replicate population. (J–M) Comparison of the numbers of COs between chromosomes in individual tetrads and each replicate population.

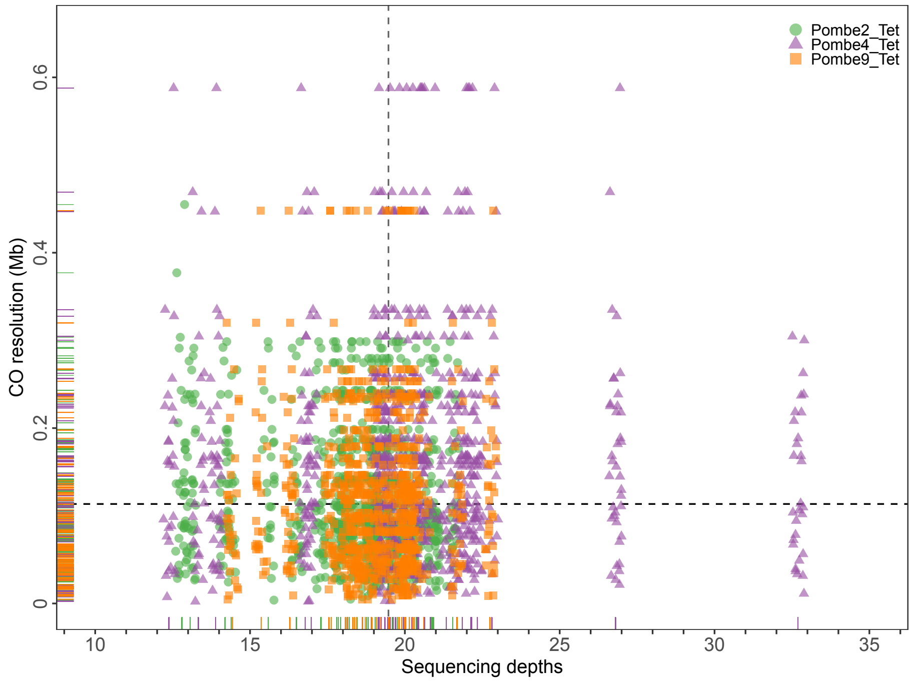


**Figure S12. Comparison of distributions of CO interval lengths and sequencing depths in each replicate population.**
The median values are indicated by dashed lines.
